# ANTIBIOFILM FORMATION ACTIVITY, RESISTANT GENES PROFILING AND DETECTION OF VIRULENCE FACTORS OF TOXIGENIC *Vibrio cholerae* ISOLATES FROM KISUMU COUNTY, KENYA

**DOI:** 10.1101/2021.01.04.425200

**Authors:** Silas O. Awuor, Omwenga O. Eric, Ibrahim I. Daud

**Affiliations:** Ministry of Health, Masogo sub-county hospital, Kisumu County P.O BOX 12-40122 Kisumu, Kenya; School of Health Sciences, Kisii University P.O BOX 408-40200 Kisii, Kenya; Kenya Medical Research Institute, United States Army Medical Research Directorate-Africa, HJF Medical Research International, Kericho, Kenya

**Keywords:** *Vibrio cholerae*, antibiofilm activity, resistant genes profiling, virulence factors

## Abstract

**Introduction:** *Vibrio cholerae* can switch between motile and biofilm lifestyles with some of its strains forming biofilms in addition to production of various virulence traits and possessing antimicrobial resistance traits. This study is aim to show antibiofilm formation activity, resistant genes profiling and detection of virulence factors of toxigenic *vibrio cholerae* isolates from Kisumu County.

**Methodology:** A total of 119 *Vibrio cholerae* O1, biotype El Tor isolates collected during 2017 cholera outbreak in Kisumu County were used for this study. The samples were cultured on TCBS and PCR assay carried out using standard procedures. Biofilm assay tests and detection of virulence factors were also done by use of standard procedures.

**Results:** Of the 101 confirmed *vibrio cholerae* isolates, 80.2% possessed the cholera toxin gene (*ctxA*) whereas 19.8% did not. Analysis of the *toxR* gene revealed that 98.0% harbored the *toxR* gene and only 2.0% did not. It was also revealed that 80.2% harbored the class I integron (*inDS* gene) while 19.8% did not, 93.1% were confirmed to possess the SXT integrating conjugative element (ICE) while 7.0% did not. The tetracycline resistance gene was present in 96.0% of the isolates. In 7 isolants strains which were resistance to common used antibiotics were screened for biofilm formation. Three of the strains (***04/17-07, 06/17-14***, and ***05/17-03***) failed to form biofilm while four strains namely ***03/17-16, 02/17-09, 04/17-13*** and *P. aeruginosa* ATCC 10145 as a positive control formed biofilms. In addition, out of those 7 isolants 71.42% produced protease, 85.71% produced phospholipases, 71.42% of isolates has the ability to produce lipase and 100% were able to produce the haemolysin.

**Conclusion:** An understanding of this intricate signaling pathway is essential for the development of methods to treat and prevent this devastating disease.

## INTRODUCTION

*Vibrio cholera* is a causative agent of cholera an acute diarrheal infection caused by ingestion of food or water contaminated with the bacterium *Vibrio cholera* that belongs to genus *vibrio*, family *Vibrionaceae* **[1]**. *Vibrio cholerae* have a number of factors which do help it to reach and colonize the epithelium of the small intestine and produce a variety of extracellular products that have deleterious effects on eukaryotic cells **[2]**.

*Vibrio cholera* have shown a number of products important for its virulence, such as cholera toxin (CT), whose action is largely responsible for the host secretory response, and toxin-co-regulated pili (TCP), which greatly enhances colonization of the intestinal epithelium **[3]**. TCP also serves as a receptor for CTXØ phage so it appears that TCP pathogenicity island is the initial genetic element required for the virulence associated genes in *V. Cholerae* **[4, 5]**. A toxin-coregulated pilus has been shown to be necessary for colonization in both the classical and El Tor biotypes of *V*.*cholerae* O1 **[6]**. Besides CT and the pilus, other factors [other potential toxins, accessory colonization factors, outer membrane proteins, proteases, hemolysins, hemagglutinins (HAs), and a capsular polysaccharide] including those which are necessary for survival of the bacteria *in vivo*, penetration of the mucous layer and adherence to the underlying epithelial cells of the intestine, binding and internalization of CT, evasion of the host defense system, etc., may also contribute to the virulence contributing to survival and multiplication of *V. cholerae* within the host **[7]**. TcpI is thought to be a chemo sensor and is proposed to be a sensor protein that negatively regulates tcpA **[5]**. ToxR dimers, but not monomers will bind to operator region of ctxAB operon and activate its transcription **[8]**. Transcription of the ctxAB operon is regulated by a number of environmental signals, including temperature, pH, osmorality and certain amino acids. Other *V. cholerae* genes are corregulated in the same manner including the tcp operon which is concerned with fimbrial synthesis and assembly. The ctx operon and the tcp are part of regulon, the expression of which is controlled by the same environmental signals **[9]**. The proteins involved in control of this regulon expression have been identified as ToxR, ToxS and ToxT. ToxR is a trans membranous protein with about two thirds of its amino terminal part exposed to the cytoplasm. ToxS is a periplasmic protein. It is thought that ToxS can respond to environmental signals, change conformation and somehow influence dimerization of ToxR which activates transcription of the operon. Expression of ToxT is activated by ToxR, while ToxT in turn activates transcription of tcp genes for synthesis of tcp pili **[10]**. Besides CT and the pilus, other factors including those which are necessary for survival of the bacteria in vivo, penetration of the mucous layer and adherence to the underlying epithelial cells of the intestine, binding and internalization of CT, evasion of the host defense system, etc., may also contribute to the virulence of this important human pathogen [**7**]. These factors included other potential toxins, accessory colonization factors, outer membrane proteins, proteases, hemolysins, haemagglutinins (HAs), and in some strains, a capsular polysaccharide, all of which may contribute to survival and multiplication of */* within the host [**6, 11**]. The genes that encode the cholera toxin subunits ctxA and ctxB are localized to a CTX genetic element which is made up of a 4.6 kbp central core region 2.4 kbp repititive sequence termed RS2. Similar RS sequences called RS1 may flank the CTX element **[12]**. Zonula occludens toxin increases the permeability of enterocytes and is encoded by *zot* gene which is within core region of CTXф **[13]**. A third toxin encoded by *ace* affects intestinal secretion **[12]**. HylA for haemolysin damages cells by acting as pore forming toxin and a number of studies have shown purified haemolysin is enterotoxic [**6**]. Outer membrane proteins which include *OmpU, OmpT, OmpS, OmpV* and others are major cell envelope proteins. *OmpU* is thought to contribute to bile resistance and functions as a colonizing factor [**14**]. Bile salts facilitate survival of *V. cholerae* in the intestine [**15**]. Toxin co-regulated pilus is clearly required for intestinal colonization [**16**].

Biofilms are. embedded in a matrix containing polysaccharides, proteins, and extracellular microbial DNA **[17, 18, 19]**. The biofilm forming pathogens like *Vibrio cholerae* and drug-resistant pathogens like MRSA cause fatal diseases to human beings and become resistant to most of the antimicrobial drugs **[20]**. This is because biofilms provides a reservoir for microbial cells, which when dispersed enhances the risk of chronic and persistent infections. It may also promote the reinfection of colonized sites **[21, 22]**. Likewise, the matrix confers a protection against biocides, Immune system activity & drugs andhas environmental promoters that induce biofilm formation that contributes to drug resistance development **[23, 24]**. All these factors contribute to biofilm cells being 1000 fold more resistant to antimicrobial agents than planktonic cells **[17]**. Equally, the current treatment and control of biofilm is complicated, because antimicrobials have been developed against planktonically grown bacteria cells and other microorganisms in metabolically active stage **[25]**.

Other virulence factors are also produced by different strains of *V. cholerae* and may be related to the hydrolysis of lipid barrier in intestinal epithelial cells **[26]**. While Neuraminidase, is secreted to cause an increase in the number of receptors in the gut **[27]**. Therefore it’s against this background that this study was to be carried out from the isolated which were found to be resistance to common used antibiotics [46].

## Materials and methods

### Study design

This was a descriptive cross-sectional study that focussed on the 2017 cholera outbreak previously isolated from diarrhoeal stool specimens. The specimens were obtained from patients presenting with passage of three or more watery stools with or without vomiting during cholera outbreaks.

### Study area

The study was performed at the six sub-counties of Kisumu County, including Muhoroni, Nyando, Nyakach, Kisumu West, Kisumu East and Seme.

### Study samples

A total of 119 isolates were sourced from KEMRI-Kisumu where they were kept following the 2017 cholera outbreak at Kisumu County. The stool samples were randomly collected from patients with severe diarrhoea who were suspected of having cholera and who were attending to different health facilities within the County in 2017 [47].

### Study procedure

#### Culturing of the V. cholerae isolates

Diarrhoeal stool samples were thawed and cultured on alkaline peptone water followed by 6 h incubation at 37°C using established procedures as used before **[28]**.The cultures were then plated on thiosulfate citrate bile salts sucrose agar (prepared using manufacturer’s guidelines). The plates were incubated at 37°C for 18–24 h, typical yellow colonies, which were presumed to be *V. cholerae* isolates were subjected to biochemical, serological and genotypic analyses using well-established protocols and as reported in our previous paper **[28]**.

#### Biofilm formation inhibition assay

As described by **[29]**, micro titter plate assay was performed to quantify the effect of commonly used antibiotics on the biofilm formation of *V. cholerae* strains. The test bacteria was first inoculated on LB agar and incubated at right conditions at 37°C overnight. Then a colony was identified, picked and inoculated in 10 mL of LB broth and incubated at 37°C overnight at 100 r.p.m for around 18 hrs. An aliquot of 190 μL of LB broth with and without antibiotic drugs (various concentrations) as used in susceptibility studies were inoculated with 10 μL of bacterial suspension per well aseptically and incubated at 37°C for 48 h without shaking. By use of a parafilm the flat-bottomed polystyrene tissue culture microplate was sealed for purposes of preventing medium evaporation. After 48hrs incubation, the wells were carefully rinsed with double-distilled water to remove loosely attached cells. The microplate was air-dried for 1 h before adding 200 μL per well of 0.4% crystal violet solution to the adhered cells in the wells and then stand at room temperature for 15 min. Excess stain was removed by rinsing gently the wells with 200 μL per well of distilled water three times. The microtiter plate was air-dried again for 1 h after which 200 μl per well of absolute ethanol was added to solubilize the dye. The Intensity was measured at OD_590_nm using a Safire Tecan-F129013 Microplate Reader (Tecan, Crailsheim, Germany). For each experiment, background staining was corrected by subtracting the crystal violet bound to untreated controls (Blank) from those of the tested sample. The experiments were done in triplicate and average OD_590_nm values were calculated. To estimate the antibiofilm activity (Abf A) of a given antibiotic the following equation was used

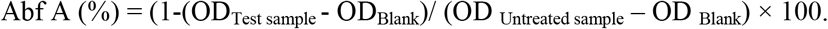

#### Molecular biology techniques/assays − PCR and Gel electrophoresis

A conventional PCR assay was performed to detect the ctxA gene for confirmation of the identification of *V. cholerae*. Other *Vibrio spp*. were screened for the presence of *CtxA, toxR, inDS, int, tetA* and *Ery* gene to determine the specificity of the assay. DNA was extracted from *V. cholerae* isolates by boiling. The specific primers selected for PCR analysis of the *ompW* gene, according to **[2]**.These primers are synthesized by the Alpha DNA Company, Canada. PCR analysis was conducted in 50 μL of a reaction mixture containing 24 μL of GoTaq Green Master, 2 μL of 25 mmol/L MgCl_2_, 2 μL of (100 pmol) primer, and 10 μL of distilled water. Amplification was conducted using a master cycler (Eppendorf) programmed with 1 cycle at 95 °C for 1 min, 40 cycles of 95 °C for 1 min, 64 °C for 1 min, 72 °C for 1 min, 72 °C for 10 min. The amplified product was subjected to 1.8% agarose gel electrophoresis, and visualized under UV after ethidium bromide staining **[4]**.

### Detection other virulence factors of V. cholerae

#### Detection of hemolysin

Haemolysin was detected according to well-established protocols as documented by **[30]**. The β-hemolytic activity was tested for on base agar (Himedia, India) supplemented with 7% sheep erythrocytes. A colony of 18-24 hr. isolates cultured on TSA (prepared according to manufacturer’s guidelines) was transferred and streaked on blood agar and incubated for 24 hr. at 37 °C. Zones of haemolysis around the colonies indicated the ability of these bacteria to haemolyse RBCs **[31]**.

#### Detection of protease

To detect protease on skim milk agar, we followed the protocol described in **[30]**. In which Solution A was prepared by adding 10 g skim milk to 90 ml of distilled water then volume was completed to 100 ml. gently heated at 50°C, then autoclaved and cooled to 50-55°C. And solution B was also prepared by adding 2 g of agar powder to 100 ml of distilled water, mixed thoroughly, then autoclaved and cooled to 50-55°C. Aseptically, 100 ml of solution A was mixed with 100 ml of solution B Then the mixture was poured into sterile petri dishes, and then stored at 4°C until to use. This media used to detect the ability of the bacteria to produce protease **[30]**. The appearance of a cleared hydrolysis zone indicates a positive test **[31]**.

#### Detection of lipase

Lipase production ability by *V. cholerae* isolates was determined by methods that have been used before [30]. Briefly a single colony of an overnight growth was cultured on Rhan medium (**Supplementary data S1**), and then incubated for 1–5 days at 37 °C. The appearance of a turbid zone around colonies indicates a positive result **[30]**.

#### Detection of lecithinase (phospholipase)

To detect lecithinase, we followed the procedure of **[32]**. In which a colony was cultured on Medium of Phospholipase Activity Assay (**Supplementary data S2**) followed by incubation for 1–3 days at 37°C using established procedures as used before **[28]**. The appearance of a white to brown color elongated precipitated zone

##### Data analysis

All experiments were conducted in triplicate to validate reproducibility. Statistical analysis was performed using Stata software. Data on graphs were analyse by Graph Pad Prism version 6. All values of diameter zones of inhibition were reported as mean ± standard error. Data analysis was performed during and after collection. The study did not involve patients.

#### Ethical consideration

Confidentiality and privacy were strictly adhered to and no names of individuals were recorded or made known in the collection or reporting of information. The study was granted ethical clearance by the Board of Postgraduate Studies (BPS) of Kisii University, and ethical approval to conduct the study was sought from the Institutional Research Ethics Committee (IREC) at Moi University/Moi Teaching and Referral Hospital (MTRH) and the National Commission of Science, Technology and Innovations (NACOSTI).

## RESULTS

### *Vibrio cholerae* O1 virulence genes

Eighty one (80.2%) isolates possessed the cholera toxin gene (*ctxA*) whereas 20 (19.8%) did not. Analysis of the *toxR* gene revealed that 99 (98.0%) harbored the *toxR* gene and only 2 (2.0%) did not. Table 1 describes the PCR results in regard to detection of pathogenic and antimicrobial resistance genes. It was also revealed that 81 (80.2%) of the isolates harbored the class I integron (encoded by *inDS* gene) while 20 (19.8%) did not. Majority, 94 (93.1%) were confirmed to possess the SXT integrating conjugative element (ICE) while 7 (7.0%) did not. The tetracycline resistance gene was present in 97 (96.0%) of the isolates. These findings add value to our previous findings on these seven (7) isolates that proved to be resistant against commonly used antibiotics in management of this condition at the study site including tetracycline that had resistance of 97.3% [46].

**Table 1:** Analysis of pathogenic and antimicrobial resistance genes (**Supplementary data S3**)by Polymerase Chain Reaction in *Vibrio cholerae* isolates from cholera outbreaks in Kisumu County, 2017 (n=101).

| Primer | Target gene | Positive n (%) | Negative n (%) |
| --- | --- | --- | --- |
| ctxA | cholera toxin | 81(80.2) | 20(19.8) |
| toxR | regulatory gene | 99(98.0) | 2(2.0) |
| inDS | class one intergron | 81(80.2) | 20(19.8) |
| int | SXT intergrase | 94(93.0) | 7(7.0) |
| tetA | tetracycline resistance | 4(4.0) | 97(96.0) |
| Ery | Erythromycin resistance | 90(90.0) | 11(10.0) |

The plates A, B, C represent agarose gel electrophoresis of DNA products resolved after virulence factors amplification.

Based on PCR analysis, *ace* gene was revealed in all the seven isolates as shown in Plate A (Figure 1). Also, *inDS, toxR* and *int* genes were revealed to be present as presented Plate B (Figure 1). Additionally, PCR genotyping also did show the presence of *ctxA* and *tcpI* genes from clinical *V. cholerae* isolates as represented in Plate C in Figure 1 above.

**Figure 1:**
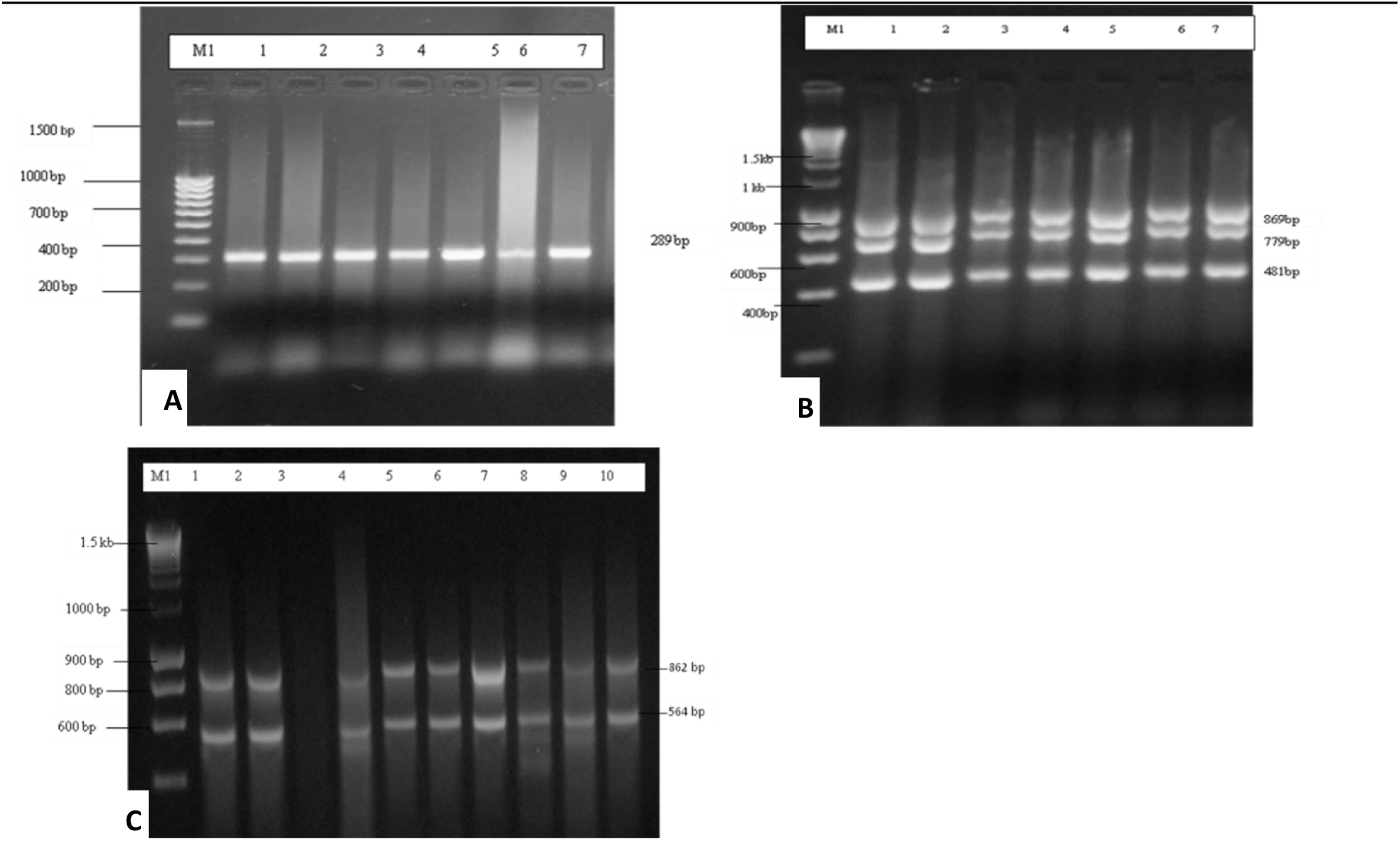
Showing the Gel electrophoresis outcome of various genes extracted from the 7 *V. cholerae* isolates that were resistant to various commonly used antibiotics with Plate A: representing the *ctxA* gene in clinical *V. cholerae*. Plate B: Representing *inDS, toxR* and *int* genes in clinical *V. cholerae* isolate and Plate C: Representing *tetA* and *Ery* genes in clinical *V. cholerae* isolates. **The legends:** Lane 1 (05/17-03 isolate^*^); Lane 2 (06/17-14 isolate^*^); Lane 3 (03/17-16 isolate^*^); Lane 4 (04/17-07 isolate^*^); Lane 5 (02/17-09 isolate^*^); Lane 6 (04/17-13Isolate^*^) and Lane 7 (06/17-07 isolate^*^). Lane M1 for Plate A represents 2 kb DNA Size Marker-Hyper ladder I *ctxA* band 289 bp; Lane M1 for Plate B represents 2 kb DNA Size Marker-Hyper ladder I *inDS* 869 bp, *toxR* 779 bp and *int* 481bp and Lane M1 for Plate B represents 2 kb DNA Size Marker-Hyper ladder I, *tetA* 862 bp and Ery 481 bp * The figures in brackets represent strain numbers.

### Biofilm formation inhibitory effects of selected antibiotics against the *V. cholerae* strains

The most resistant drugs towards the seven isolants (Tetracycline, Ampicillin, Amoxicillin, Cotrimoxazole, Erythromycin and Nalidixic Acid) *were* subjected to biofilm formation inhibition assay in a 96-well microtiter plate. Different concentrations of the drugs were prepared using two-fold serial dilution (with dosage ranging from 0.5 to 0.03125mg/ml) and the wells were inoculated with 10 μL of ***V. cholerae*** isolants. *P. aeruginosa* served as the positive control. The results showed that the biofilm formation inhibitory effects of the various concentrations (0.5mg/mL, 0.25 mg/ml, 0.125mg/ml, and 0.0625 mg/dl up to 0.03125 mg/mL) were significantly lower than that of the positive control, an indication that biofilm formation was inhibited at these concentrations as shown in **figures 2** to **7 below**. As much as such inhibitory effects were recorded these findings clearly demonstrate that the 7 isolates that proved to be resistant to commonly used antibiotics do form biofilms.

**Figure 2:**
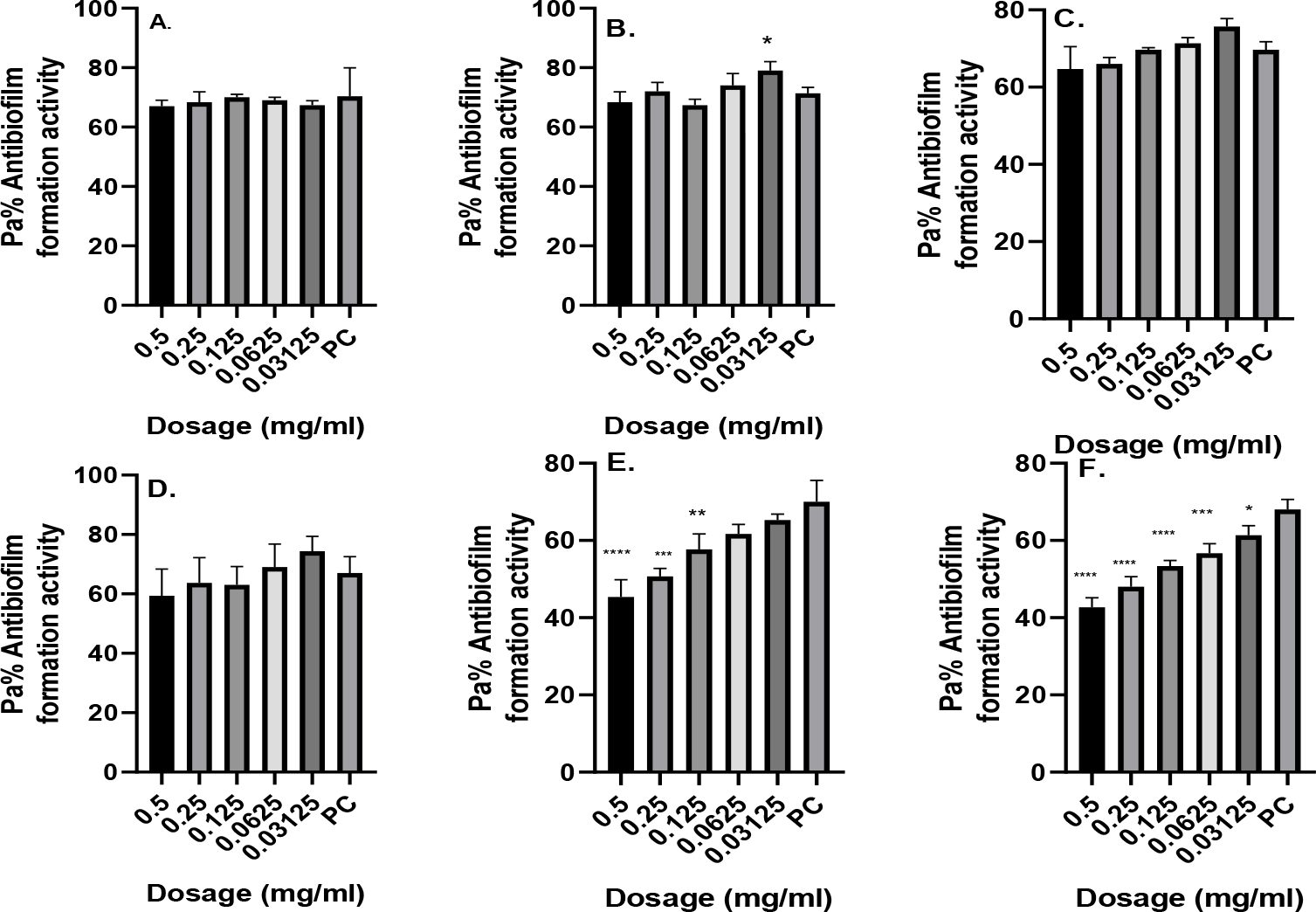
Antibiofilm formation activity against isolate **03/17-16** of *V. cholerae* against various antibiotics **A**. Tetracycline **B**. Ampicillin, **C**. Amoxicillin, **D**. Cotrimoxazole, **E**. Erythromycin & **F**. Nalidixic Acid; PC= *P. aeruginosa*-Positive control (n=3, ANOVA Dunnett’s multiple comparisons test; * P = 0.05; ** P=0.01; *** P=0.001; **** P=0.0001).

In **Figure 2** antibiofilm formation activity against isolate 03/17-16 of *V. cholerae* against various antibiotics like Tetracycline, Ampicillin, Amoxicillin, Cotrimoxazole, Erythromycin & Nalidixic Acid was observed in all the antibiotics. On the ampicillin treatment at the concentration of 0.03125 mg/mL yielded significant biofilm inhibition as compared with the positive control (p = 0.05), while for erythromycin their significant biofilm formation inhibition at a concentration of 0.5 mg/mL (p = 0.0001), 0.25mg/mL (p= 0.001), 0.125 mg/mL (p =0.01) as compared with the positive control. Nalidixic Acid treatment yielded significant biofilm inhibition at concentration of 0.5 mg/mL, 0.25 mg/mL, 0.125 mg/mL, 0.0625 mg/mL (p = 0.0001) and 0.03125 mg/mL (p = 0.05) as compared with the positive control. It is more worrying that tetracycline a commonly used antibiotic in management of the cholera at the study site had less inhibitory effects on biofilm formation, a fact that also supports our previous findings where we did document that the isolates were resistant to this antibiotic.

In **Figure 3** antibiofilm formation activity against isolate 02/17-09 of *V. cholerae* against various antibiotics was seen in all the antibiotics. All the antibiotics showed some significance difference at various dosages as compared with the positive control (**Figure 3 above**). Generally amoxicillin did not have good inhibitory effects against this isolate. On the other hand Cotrimoxazole showed a reverse activity with high inhibitory effects being observed at low dosages as compared with higher concentrations. In comparison with the positive control isolate, there was more inhibitory effects against the test isolate as compared to it.

**Figure 3:**
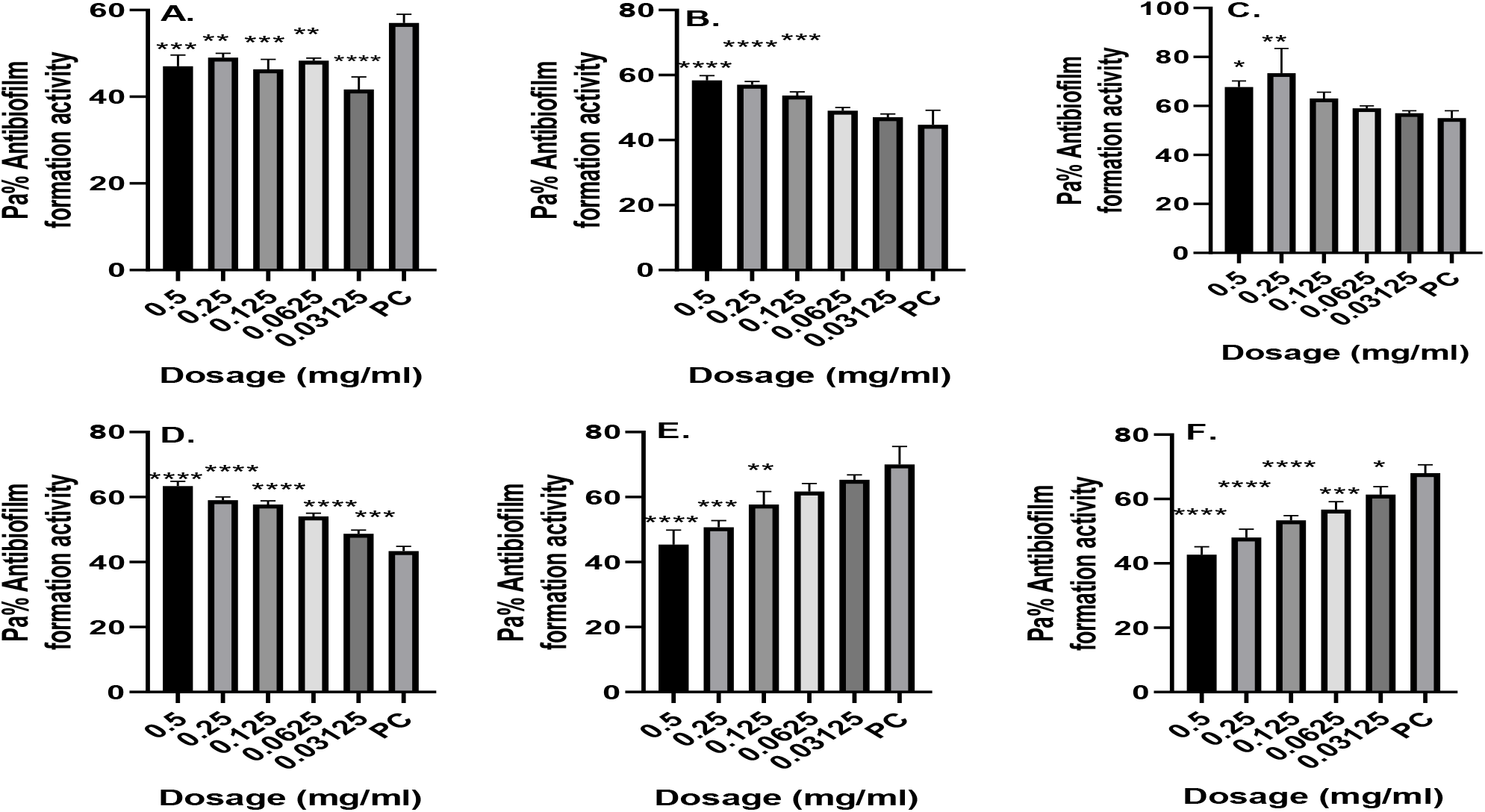
Antibiofilm formation activity against isolate **02/17-09** of *V. cholerae* against various antibiotics **A**. Tetracycline **B**. Ampicillin, **C**. Amoxicillin, **D**. Cotrimoxazole, **E**. Erythromycin & **F**. Nalidixic Acid; PC= *P. aeruginosa*-Positive control (n=3, Anova Dunnett’s multiple comparisons test; * P = 0.05; ** P=0.01; *** P=0.001; **** P=0.0001)

In **Figure 4** antibiofilm formation activity against isolate 04/17-13 of *V. cholerae* against various antibiotics was seen in all the antibiotics with most antibiotics producing significant differences as compared with the positive control at various dosages as shown in **figure 4** above. With this test isolate also inhibitory activities were observed more at lower dosages as compared at high dosages in all antibiotics bioassyed.

**Figure 4:**
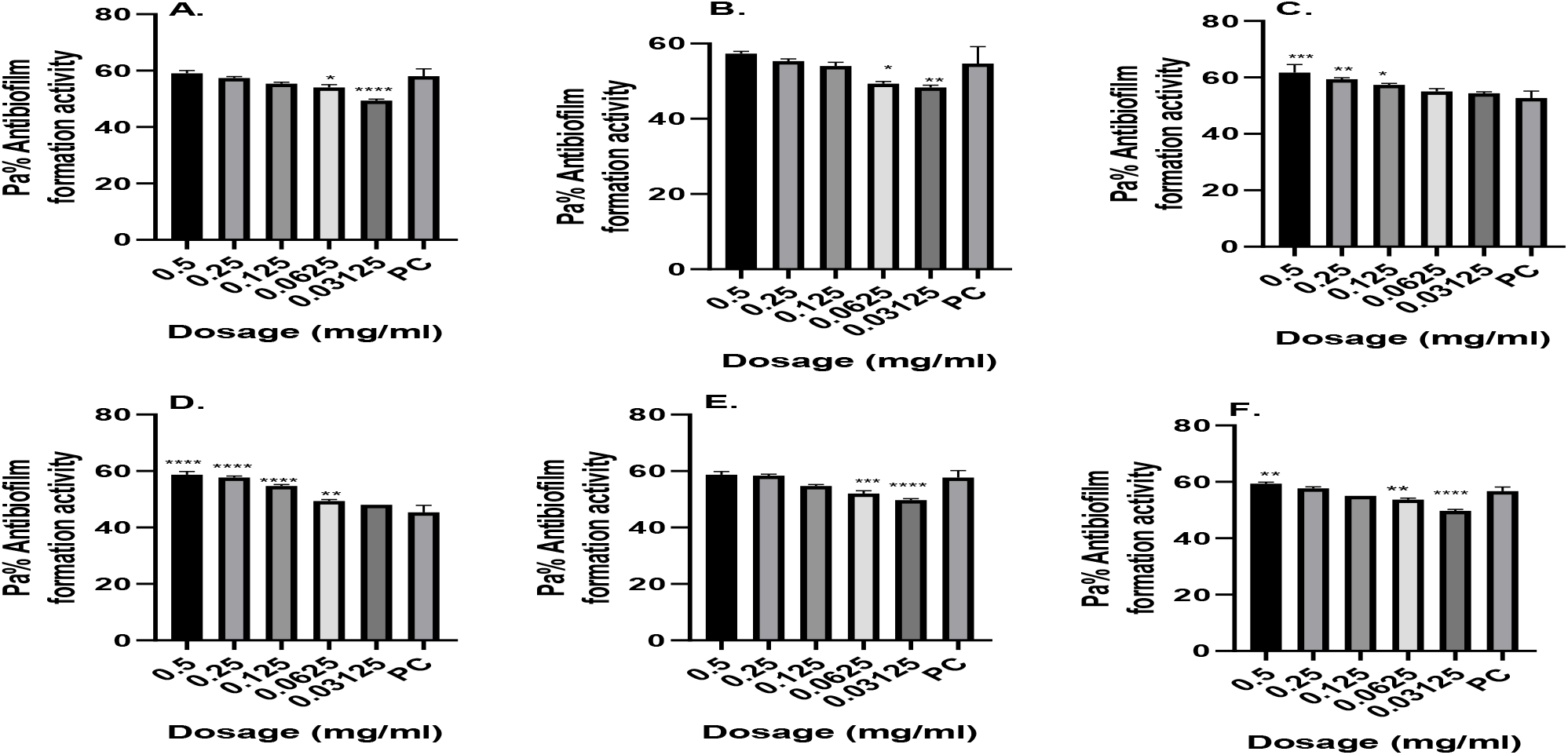
Antibiofilm formation activity against isolate **04/17-13** of *V. cholerae* against various antibiotics **A**. Tetracycline **B**. Ampicillin, **C**. Amoxicillin, **D**. Cotrimoxazole, **E**. Erythromycin & **F**. Nalidixic Acid; PC= *P. aeruginosa*-Positive control (n=3, Anova Dunnett’s multiple comparisons test; * P = 0.05; ** P=0.01; *** P=0.001; **** P=0.0001)

This test isolate was more prone to the inhibitory effects of the various dosages of the various antibiotics as compared to other isolates. Significance difference was also observed almost in each antibiotic used as shown in Figure 5 above when they were compared with the positive control.

**Figure 5:**
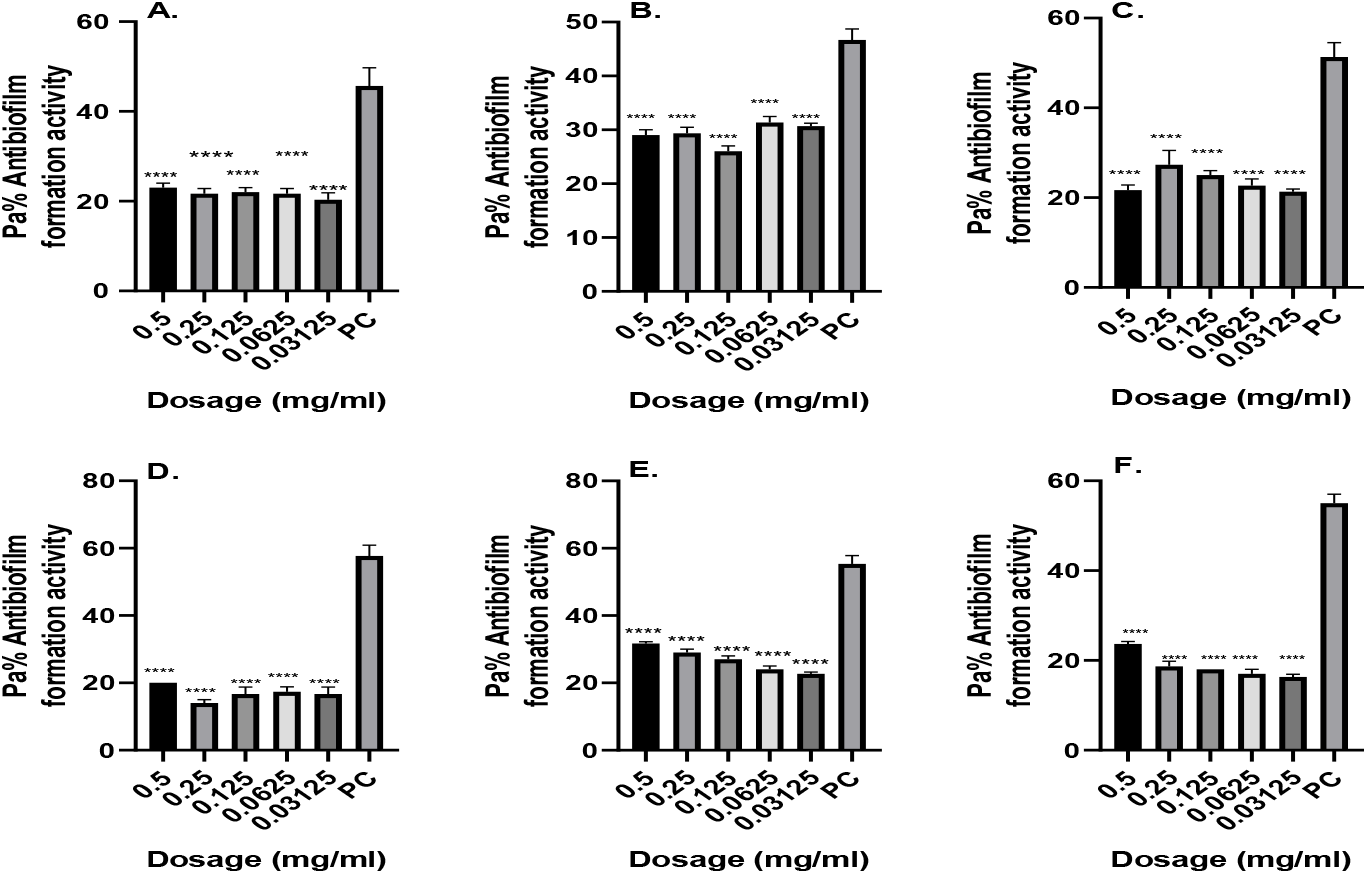
Antibiofilm formation activity against isolate **05/17-07** of *V. cholerae* against various antibiotics **A**. Tetracycline **B**. Ampicillin, **C**. Amoxicillin, **D**. Cotrimoxazole, **E**. Erythromycin & **F**. Nalidixic Acid; PC= *P. aeruginosa*-Positive control (n=3, Anova Dunnett’s multiple comparisons test; * P = 0.05; ** P=0.01; *** P=0.001; **** P=0.0001)

In **Figure 6** antibiofilm formation activity against isolate **05/17-03** of *V. cholerae* against various antibiotics was also reduced in all the antibiotics. Interestingly, the concentrations of 0.5 mg/mL, 0.25 mg/mL, 0.125 mg/mL, 0.0625 mg/mL and 0.03125 mg/mL were able to inhibit biofilm formation much more as compared with the positive control with a significance differences observed at most dosages as represented in **Figure 6** above.

**Figure 6:**
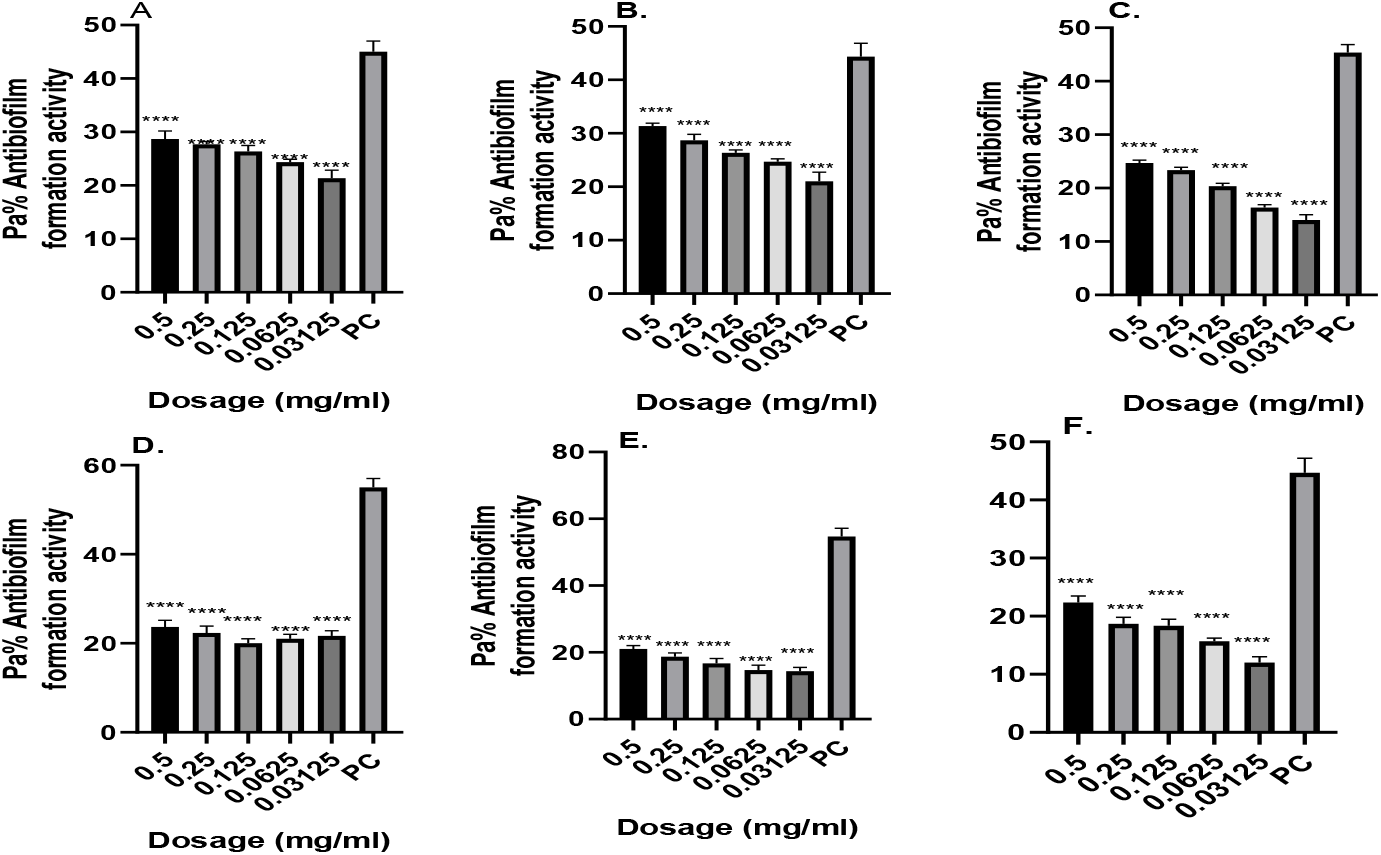
Antibiofilm formation activity against isolate **05/17-03** of *V. cholerae* against various antibiotics **A**. Tetracycline **B**. Ampicillin, **C**. Amoxicillin, **D**. Cotrimoxazole, **E**. Erythromycin & **F**.Nalidixic Acid; PC= *P. aeruginosa*-Positive control (n=3, Anova Dunnett’s multiple comparisons test; * P = 0.05; ** P=0.01; *** P=0.001; **** P=0.0001)

In **Figure 7** antibiofilm formation activity against isolate **06/17-14** of *V. cholerae* against various antibiotics was seen in all the antibiotics with no significance difference on tetracycline at all the concentration as compared with the positive control. The rest of the antibiotics produced significance differences at various dosages as compared with the positive control (see **figure 7** above). Erythromycin, tetracycline and Nalidixic Acid also showed much more inhibitory effects against this isolate at lower dosage.

**Figure 7:**
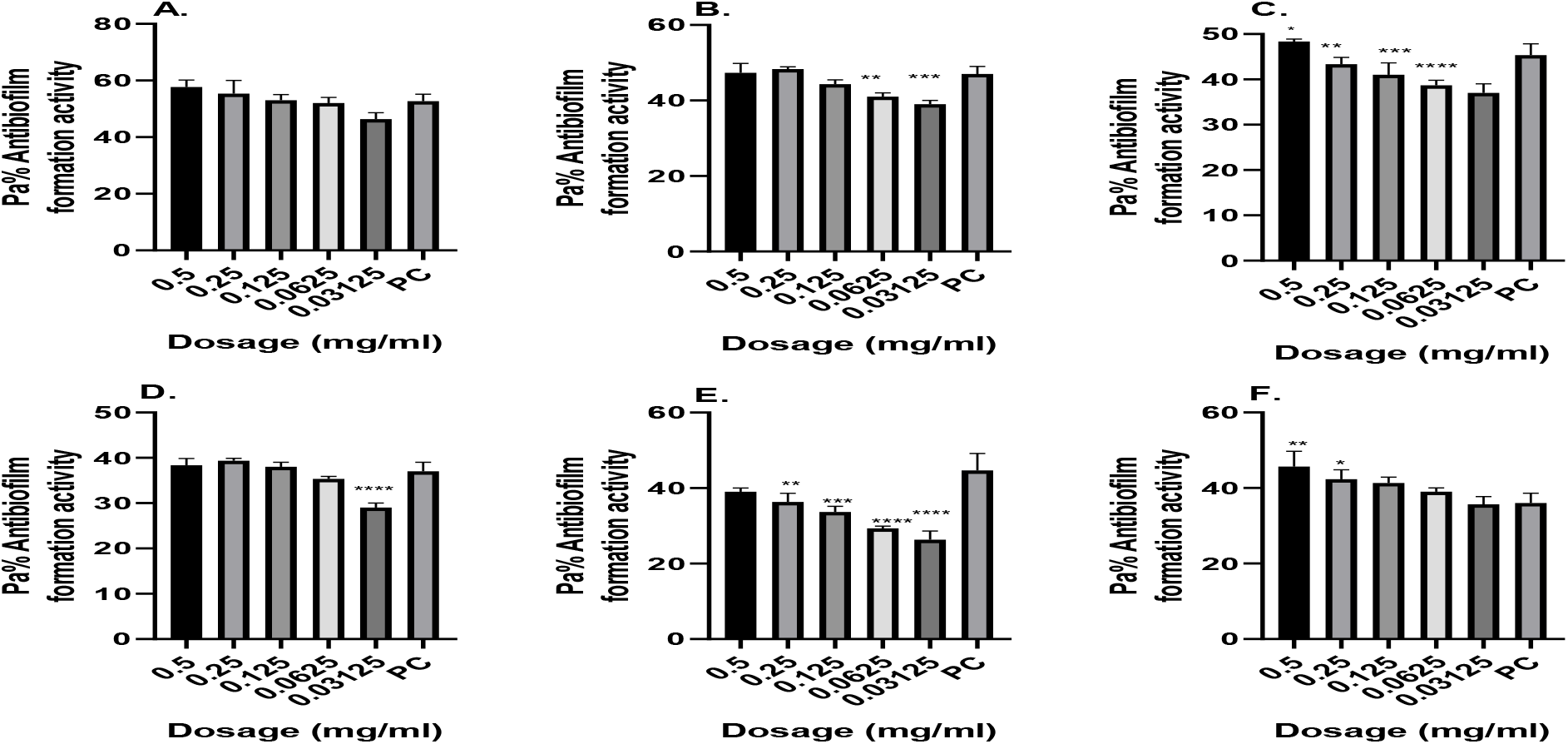
Antibiofilm formation activity against isolate **06/17-14** of V. cholerae against various antibiotics **A**. Tetracycline **B**. Ampicillin, **C**. Amoxicillin, **D**. Cotrimoxazole, **E**. Erythromycin & **F**. Nalidixic Acid; PC= *P. aeruginosa*-Positive control (n=3, Anova Dunnett’s multiple comparisons test; * P = 0.05; ** P=0.01; *** P=0.001; **** P=0.0001)

### Detection of Some Virulence Factors of *V. cholerae*

In this study the production of various virulence enzymes like protease, phospholipase, lipase and haemolysin were studied on the 7 isolates which were found to be resistant to all drugs examined from our previous study. It seemed that 5/7 (71.42%) of these isolates of *V*.*cholerae* produced protease enzyme (**Fig.8-plate A**). Also it was deduced that 6 out 7 isolates (85.71%) produce phospholipases (**Fig.8– plate B**). Further findings also indicate that out of the 7 isolates 5 isolates (71.42%) had the ability to produce lipase (**Fig.8– plate D**). Lastly, it was also deduced that all of the 7 isolates (100%) were able to produce the haemolysin by haemolysing the sheep red blood cells causing beta (β) haemolysis as shown in (**Fig.8– plate C**) above. Haemolysin therefore was the most produced virulence factor by these isolates (100%) and it was followed by phospholipase (85.71%) and lastly lipase and protease (71.42%). Out of the seven isolates 4 isolates (03/17-16, 02/17-09, 04/17-13 and 06/17-07) produced all the virulence traits studied, 2 isolates (06/17-14 and 05/17-07) produced at least 3 virulence traits and 1 isolate (05/17-03) produced at least one of the virulence traits.

**Figure 8:**
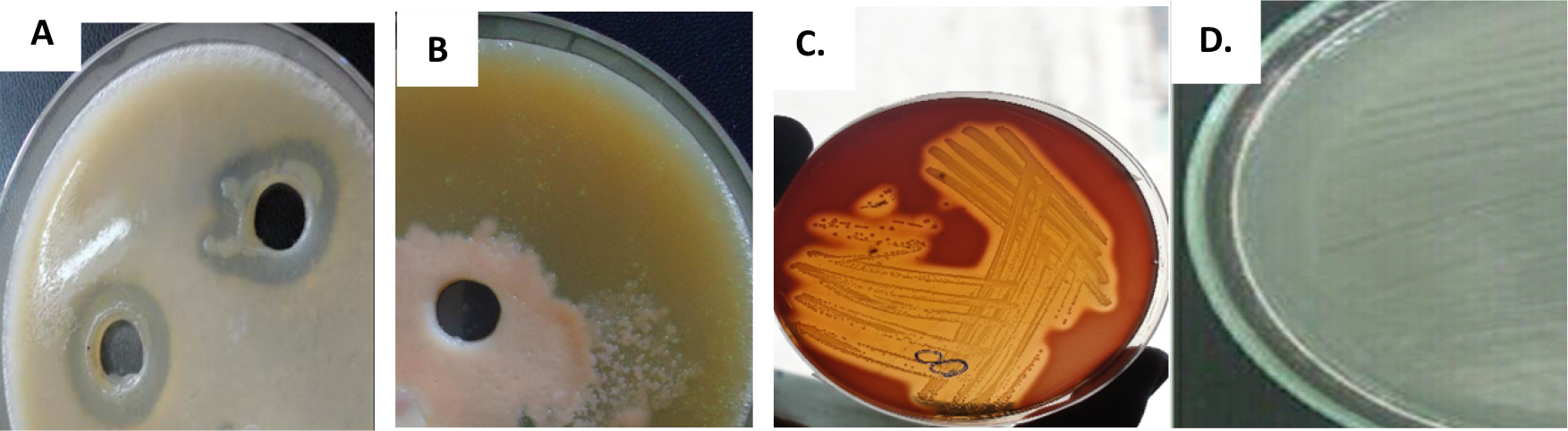
Representation of the various virulence traits produced by the *V. cholerae* isolates, **A**. The distribution of Protease enzyme production of *V. cholerae*, **B**. The Phospholipase enzyme production of *V*.*cholerae*, **C**. The Haemolysin enzyme production of *V*.*cholerae –* haemolytic activity, **D**. Lipase enzyme production of *V*.*cholerae*.

## DISCUSSION

In this study it’s clear that the seven isolates that had been discovered from our previous study to be resistant to various commonly used antibiotics in deed possess various factors that enable them either to be resistant or induce various infections in human kind. Some of these traits are controlled at the gene level and as such they can be passed on from one bacteria cell to another [46] Antimicrobial drug resistance in *Vibrio* species for instance, may arise through mutation or through acquisition of resistance genes on mobile genetic elements like plasmids, transposons integrons, and integrating conjugative elements **[33]**. Isolates analysed in this study possessed the class I integron and the SXT integrating conjugative element. Genetic elements like the class I integron (*inDS*) and the integrating conjugative elements such as SXT have been associated with the spread of genetic determinants, encoding for antimicrobial resistance in *Vibrio cholerae* **[34]**. The SXT element has been reported to harbor genes encoding for resistance to chloramphenicol (encoded by *floR*), streptomycin (encoded by *strA and strB*), trimethoprim (encoded by *dfrA18*) and sulfamethoxazole (encoded by *sul2*) **[35]**. The class 1 integron has also been reported to harbor aminoglycoside resistant gene cassettes in *Vibrio cholerae* O1 isolates cholerae **[36]**. As such therefore, resistance to erythromycin in the analysed isolates could be attributed to the presence of the class 1 integron gene that was found to be present in some strains isolated.

The finding of susceptible isolates towards streptomycin but still amplifying a 383 bp fragment of the *strA* gene suggests that this gene is not an intrinsic feature of this family of integrase, but rather appears to have been inserted into these elements, becoming transmissible in bacterial populations, as reported by other investigators **[37]**. Nalidixic acid resistance observed in this study could be attributed to mutations in the *gyrA* gene. Many investigators have reported *gyrA* gene mutations in fluoroquinolones resistant clinical isolates of *Vibrio cholerae* since it contains the active site tyrosine that forms a transient covalent intermediate with DNA hence making it to be resistance to the drug **[38]**. However, more study needs to be done to confirm this.

The results also suggest that erythromycin and Nalidixic Acid do inhibit growth at high concentrations but from our study, it was deduced that they inhibit biofilm formation at lower concentrations. Same scenario was also observed in most antibiotics like tetracycline, Amoxicillin, Cotrimoxazole etc with such “Goldilocks” effect. A possible explanation to the less activity observed at greater doses could be associated to the aggregation effects of the antibiotics at site of entry into bacteria cell especially at high dosages something that is not observed at lower dosages. It is likely that aggregation may favour biofilm formation as antibiotics struggle to reach at the point of action and hence bacteria will continue to thrive hence form more biofilms **[39]**. This finding agrees with the previous studies on biofilm inhibition by Taganna which show higher biofilm inhibitory at lower dosage concentration against the positive control **[40]**. However, it did not concur with the previous study on biofilm inhibition by Vasavi (2013) which shows that Erythromycin growth inhibitory at high concentration but also biofilm inhibitory at high concentration as compared with positive control **[41]**. On the other hand, Tetracycline, Ampicillin, Amoxicillin and Cotrimoxazole were found to inhibit biofilm formation at higher concentration in some isolates and against the positive control. However, it should be noted that they did not fully inhibit biofilm formation ability of the test isolates a clear indication that proper antibiotics should be used in management of conditions caused by these isolates. Our finding also agrees with the previous study on biofilm inhibition by Vasavi which shows that most common antibiotic used in *V. cholerae* management have a growth and biofilm inhibitory at high concentration as compared with positive control **[41]**.

However, isolates 05/17-07 and 05/17-03 of *V. cholerae* seemed to be very sensitive to the antibiotics screened at various dosages. This clearly demonstrates that the biofilm formation by these isolate can easily be managed by these antibiotics. Since they proved to be resistant when they were subjected to antimicrobial tests from our previous study it’s clear that they could be possessing other mechanisms of activity against the antibiotics screened. One of such possible mechanisms could be presence of multi drug efflux pumps presence in them as they have been found to be present in some strains of *V. cholerae* **[30]**. This make the bacteria to be resistance to antibacterial agents and other toxic compounds by a mechanism known as active efflux, where the integral membrane transporters known as drug efflux pumps, prevent the accumulation of drugs inside the bacterial cells**[40]**.

In addition to antibiofilm formation determination we also investigated the ability of these test strains to produce various virulence factors which may play role in their pathogenicity. Among the virulence traits we examined include detection of proteases, lipases, haemolysin and phospholipase. We did find that (71.42%) of the isolates of *V. cholerae* produced protease. This finding concurs to the findings of a previous study done by **[42]**, which shows that (72.7%) of isolates were protease positive and that this enzyme have limited effect on the pathogenesis of this bacteria. Our findings also concurred with **[10]** study that documented that all isolates in her study were having ability to protease production. Proteases produced by *V*.*cholerae* have a very important role in its pathogenicity, due to the hydrolysis of several physiologically important protein such as mucin, fibronectin, and lactoferrin **[43]**. It could also proteolytically activate cholera toxin A subunit, El Tor cytolycin and haemolysin hence making this pathogen to be more virulent **[44]**.

For phospholipases, 85.71% of the isolates were found to be positive. These findings do concur with the previous study findings that were done by **[9]**, who did report that out of 20 isolates 13 (67%) of isolates were having the ability to phospholipase production. Our findings are also in tandem with **[45]** study findings that documented that all of his isolates were phospholipase positive. [36] mentioned the role of this enzyme in the cholera disease by the release of Arachidonic acid from the phospholipid found in the cell membranes of the lumen cells, this plays an important role in the prostaglandin E2 (PGE2) production which is responsible for the increase of liquids secretion from the lumen cells and this lead to watery **[36]**. Therefore, its presence makes *V. cholerae* isolate to be more virulent in watery diarrhoea production a key symptom of cholera.

Also, out of the 7 isolates 5 (71.42%) had the ability to produce lipase. Our findings also concur with other studies **[10]**, which showed that all isolates in her study were having the ability to lipase production. Abbas, (2006) Abass also did demonstrate that 65.4% of his isolates had the ability to produce lipases **[42]**. Lipases enzymes catalyse the hydrolysis of the ester bonds of triacylglycerols and may have a critical role in *V*.*cholerae* pathogenicity or nutrition acquisition. The production of excesses amount of lipases allows bacteria to penetrate fatty tissue with the consequent formation of abscesses **[3]**. The production of these enzymes by the isolates may reflect the presence of genetic organization of a discrete genetic element which encodes three genes responsible for the production of proteases, lipases and phospholipase. This organization could be a possible part of pathogenic island, encoding a product capable of damaging host cells and being involved in nutrient acquisition **[43]**.

In this study 100% of, the isolants were able to produce the haemolysin. A finding that concurs to the findings of a previous study done by Abbass in which 100% of isolates were haemolysin positive. Halpern and Izhaki stated that the purified haemolysin is capable of causing fluid accumulation, in contrast to the watery fluid produced in response to CT, the accumulated fluid produced in response to haemolysin with invariably bloody with mucous **[1]**.

## Conclusion

From this study, we can conclude that most clinical isolates produced virulence factors such as heamolysin, lipase, protease and phospholipase. As much as inhibitory effects were recorded, these findings clearly demonstrate that the isolates proved to be resistant to commonly used antibiotics and they do form biofilms. and these findings therefore, add value to our previous findings on these seven isolates that proved to be resistant against commonly used antibiotics in management of this condition at the study site. To the best of our knowledge, this is the first time such data has been documented from the study site that has had a good number of cholera outbreaks before and we hope it will assist in management of this condition.

## Acknowledgement

We wish to sincerely acknowledge and thank KEMRI-Kisumu, Kenya for proving us with the Stored samples of 2017 cholera outbreak in the study area. We also thank AMPATH Laboratories– Eldoret, Kenya for providing us the bench space and reagents that made this study a success and Elsy jebitok the research assistant who assisted on the lab work.

## Authors’ contributions

All authors contributed equally to this work.

## Data availability statement

Data sharing is not applicable to this article as no new data were created or analysed in this study.

## Disclaimer

The views and opinions expressed in this article are those of the authors and do not necessarily reflect the official policy or position of any affiliated agency of the authors

## Supplementary data

**Supplementary data S1: Rhan Media of Lipase Activity Assay preparation**

It was prepared by dissolving 5 g of K_2_HP0_4_, 5 g of (NH_4_)_2_PO_4_, 1 g of Cacl2. 6H_2_O, 1g of MgSO_4_. 7H_2_O, 0.001 g of Fecl_2_.6H_2_O, 0.001 g of NaCl, 20 g of agar powder and 5 ml of olive oil in 900 ml of distilled water. Then the volume was completed to 1000 ml, final pH was adjusted to 7.2, then autoclaved, cooled and poured in sterile Petri dishes and stored at 4°C until to use. This media used to detect the ability of the bacteria to produce lipase (Rodina et al., 2018).

**Supplementary data S2: Medium of Phospholipase Activity Assay**

Prepared by dissolving 2.4 g of nutrient agar in 100 ml of distilled water with 1 g of NaCl. After autoclaved and cooled to 50°C, the addition egg yolk of one egg was done in a septical condition. Then mixed well and poured into sterile Petri dishes and stored at 4°C until to be use between 24-48 hours (Dogan et al., 2003).

**Supplementary data S3:**
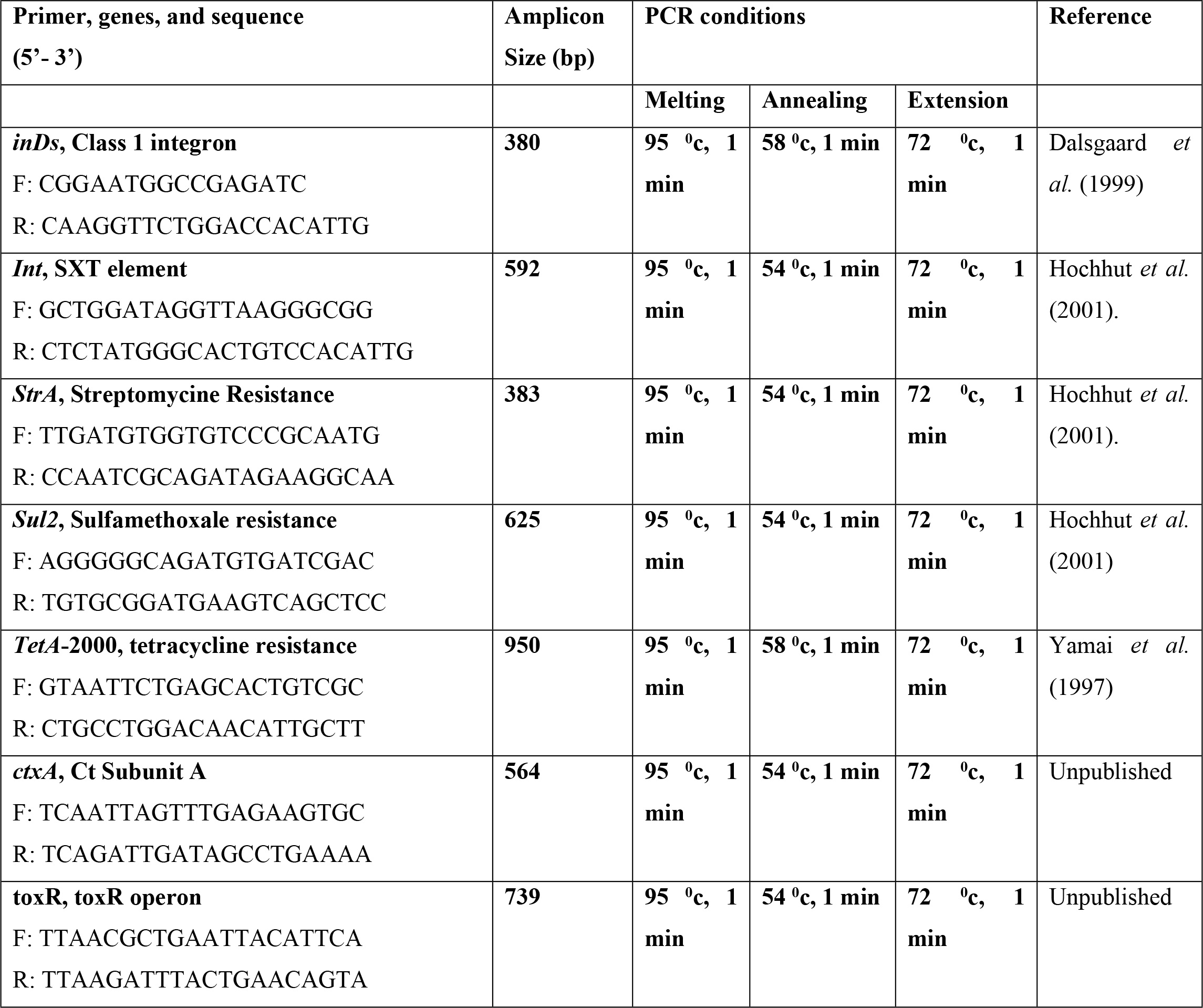
Oligonucleotide primers, sequences, amplicons and conditions used in PCR assays in this study.

## REFERENCE

1. Halpern, M and Izhaki I. Fish as Hosts of Vibrio cholerae. Frontiers in Microbiology. 2017; 8:282.doi: 10.3389/fmicb.2017.00282.

2. Matthey, N, and Blokesch, M. “The DNA-Uptake Process of Naturally Competent Vibrio cholerae”. Trends Microbial. 2016; 24 (2): 98–110. PMID 26614677. doi:10.1016/j.tim.2015.10.008.

3. Sakib SN, Reddi G, and Almagro-Moreno S. Vibrio JB Special Issue. Minireview Environmental role of pathogenic traits in Vibrio cholerae.J. Bacteriol.2018; doi:10.1128/JB.0079517. American Society for Microbiology.

4. Siriphap A, Leekitcharoenphon P, Kaas RS, Theethakaew C, Aarestrup FM, Sutheinkul O, et al. Characterization and Genetic Variation of Vibrio cholerae Isolated from Clinical and Environmental Sources in Thailand PLoS ONE.2017;12(1):e0169324. https://doi.org/10.1371/journal.pone.0169324.

5. AL-Fatlawy HNK, Aldahhan HA, and Alsaadi AH. Phylogenetic of DNA Fingerprinting and New Sequencing of Aeromonas Species and V. Cholerae DNA. American Journal of Applied Sciences 2017; 14 (10): 955.964. DOI: 10.3844/ajassp.2017.955.964.

6. Moore S, Thomson N, Mutreja A, Piarroux R. Widespread epidemic cholera caused by a restricted subset of Vibrio cholerae clones. Clin Microbiol Infect. 2014; 20(5):373–9. Pmid: 24575898.

7. Kirn TJ, Lafferty MJ, Sandoe CM, Taylor RK. Delineation of pilin domains required for bacterial association into microcolonies and intestinal colonization by Vibrio cholerae. Mol Microbiol. 2000; 35 (4):896–910. Pmid: 10692166.

8. Al-Hadrawy HAN. A Comparative study of Bacteriological and Molecular Vibrio Cholera Isolated from the Tigris and Euphrates. Ph. D. Thesis, College of the science, University of Kufa in Arabic 2012.

9. Jabik NA. Study of some Genetic Aspects of Isolated V. cholerae in Babylon. M.Sc. Thesis, College of Science, University of Babylon 2000.

10. Al-Khafaji KAA. Identification of Some Virulence Factors in Toxigenic Clinical and Environmental Isolates of Vibrio cholerae. MSc. Thesis. Genetic Engineering and Biotechnology. University of Baghdad 2007.

11. AL-Fatlawy HNK and Al-Ammar MH. Study of Some Virulence Factors of Aeromonas Hydrophila Isolated from Clinical Samples (Iraq). International Journal of Science and Engineering Investigations, Volume 2, Issue 21, October 2013. Paper ID: 22113-16. (2013). WHO. (2002).

12. Jawetz E, JI Melnick and EA Adelberg. Medical Microbiology. 27th Edn. Appleton and Lange U.S.A. 2016.

13. Yang QH, Zhou C, Lin Q, Lu Z, He LB, Guo SL. Draft genome sequence of Aeromonas sobria strain 08005, isolated from sick Rana catesbeiana 2017 Genome Announc 5:e01352–16. https://doi.org/10.1128/genomeA.01352-16.

14. Provenzano D, DA Schuhmacher, JL. Barker and KE Klose. The Virulence Regulatory Protein ToxR Mediates Enhanced Bile Resistance in Vibrio cholerae and Other Pathogenic Vibrio Species 2000. Infection and Immunity 68:1491–1497.

15. Fasano A. Bacterial infections: small intestine and colon. Current opinion in Gastroenterology 17:4–9 2001.

16. Reguera G and R Kolter. Virulence and the environment: a novel role for Vibrio cholerae toxin-coregulated pili in biofilm formation on chitin 2005. Journal of Bacteriology 187:3551–5.

17. Taraszkiewicz A, Fila G, Grinholc M, Nakonieczna J. Innovative strategies to overcome biofilm resistance 2013. Biomed Res Int 2013: 150653.

18. Simões M, Simões LC, Vieira MJ. A review of current and emergent biofilm control strategies 2010. LWT-Food Science and Technology 43: 573–583.

19. Lazar V. Quorum sensing in biofilms--how to destroy the bacterial citadels or their cohesion/power? Anaerobe 2011. 17: 280–285.

20. Faruque SM, Biswas K., Udden SM, Ahmad QS, Sack DA, Nair GB, et al. Toxin, toxin-coregulated pili, and the toxR regulon are essential for Vibrio cholerae pathogenesis in humans 2006. J. Exp. Med. 168: 1487–1492.

21. Høiby N, Bjarnsholt T, Givskov M, Molin S, Ciofu O. Antibiotic resistance of bacterial biofilms 2010. Int J Antimicrob Agents 35: 322–332.

22. Chen L, Wen YM. The role of bacterial biofilm in persistent infections and control strategies 2011. Int J Oral Sci 3: 66–73.

23. Tan SY, Chew SC, Tan SY2, Givskov M3, Yang L4. Emerging frontiers in detection and control of bacterial biofilms 2014. Curr Opin Biotechnol 26: 1–6.

24. Poole K. Bacterial stress responses as determinants of antimicrobial resistance 2012. J Antimicrob Chemother 67: 2069–2089.

25. Ammons MC. Anti-biofilm strategies and the need for innovations in wound care 2010. Recent Pat Antiinfect Drug Discov 5: 10–17.

26. Pride AC, Guan Z and Trent MS. Characterization of the Vibrio cholerae VolA Surface-Exposed Lipoprotein Lysophospholipase 2014. J. Bacteriol. 2014, 196(8):1619. DOI: 10.1128/JB.01281-13.

27. AL-Fatlawy HNK and Al-Ammar MH. Study of Some Virulence Factors of Aeromonas Hydrophila Isolated from Clinical Samples (Iraq) 2013. International Journal of Science and Engineering Investigations, Volume 2, Issue 21, October 2013. Paper ID: 22113–16.

28. CLSI. Performance standards for antimicrobial sensitivity testing. Twentieth Informational Supplement 2010. M100-S20. Volume 30.

29. Omwenga EO, Hensel A, Pereira S, Shitandi AA, & Goycoolea FM. Antiquorum sensing, antibiofilm formation and cytotoxicity activity of commonly used medicinal plants by inhabitants of Borabu sub-county, Nyamira County, Kenya 2017. PLoS one, 12(11), e0185722.

30. Benson HJ. Microbiological Applications: Laboratory Manual in General Microbiology. (8th edit). Complete version. McGraw-Hill. U.S.A. 2002.

31. Elliot EL, Kaysner CA, Jackson L and Tamplin ML. V. cholerae, V. parahemolyticus, V. Valnificus and other Vibrio spp. In: Food and Drug Administration: Bacteriological Analytical Manual, chapter 9, 8th ed. edited by Merker, R. L., AOAC International, Gaithersburg, MD 2001.

32. Dogan B and Boor KJ. Genetic diversity and spoilage potentials among Pseudomonas spp. Isolated from fluid milk products and dairy processing plants 2003. Appl. Environ. Microbiol. 69 (1): 130–138.

33. Sjölund-Karlsson M, Reimer A, Folster J P, Walker M, Dahourou G A, Batra D G, Martin I, Joyce K, Parsons MB & Boncy J. Drug resistance mechanisms in Vibrio cholerae O1 outbreak strain 2011. Emerging Infectious Diseases, 17, 2151–4.

34. Dalsgaard A, Forslund A, Sandvang D, Arntzen L & Keddy K. Vibrio cholerae O1 outbreak isolates in Mozambique and South Africa in 1998 are multiple-drug resistant; contain the SXT element and the aadA2 gene located on class 1 integrons 2001. Journal of Antimicrobial Chemotherapy, 48, 827.

35. Beaber JW, Hochhut B & Waldor MK. Genomic and functional analyses of SXT, an integrating antibiotic resistance gene transfer element derived from Vibrio cholerae 2002. Journal of bacteriology, 184, 4259.

36. Oliver J D and Kaper JB. Vibrio Species. In: Food Microbiology: Fundamentals and Frontiers. edited by Doyl, MP, Beuchat LR and Montville TJ, ASM press, Washington D C, USA 2007. Pp. 228–60.

37. Hochhut B, Lotfi Y, Mazel D, Faruque SM, Woodgate R & Waldor MK. Molecular analysis of antibiotic resistance gene clusters in Vibrio cholerae O139 and O1 SXT constins 2001. Antimicrobial Agents and Chemotherapy, 45, 2991–3000.

38. Baranwal S, Dey K, Ramamurthy T, Nair GB & Kundu M. Role of active efflux in association with target gene mutations in fluoroquinolones resistance in clinical isolates of Vibrio cholerae 2002. Antimicrobial agents and chemotherapy, 46, 2676.

39. Xu KD, McFeters GA. & Stewart PS. Biofilm resistance to antimicrobial agents 2000. Microbiology. 146, 547–549.

40. Taganna J, Quanico J, Perono R, Amor E, Rivera W. Tanninrich fraction from Terminalia catappa inhibits quorum sensing (QS) in Chromobacterium violaceum and the QS-controlled biofilm maturation and LasA staphylolytic activity in Pseudomonas aeruginosa2011. J. Ethnopharm 134(3): 865–871.

41. Vasavi HS, Arun AB, Rekha PD. Inhibition of quorum sensing in Chromobacterium violaceum by Syzygium cumini L. and Pimenta dioica L. Asian Pac. J. Trop. Biomed. 2013; 3(12), 954–959. Doi: 10.1016/S22211691(13)60185-9.

42. Abbass NBM. Effectiveness of some physical and chemical factors on the morphological changes of Vibrio cholerae isolated from environment. Ph.D. Thesis, College of Science, University of Al Mustansyria 2006.

43. Namdari H, Klaips CR and Hughes JL. A cytotoxin-producing strain of Vibrio cholerae Non-O1, Non-O139 as a cause of cholera and bacteremia after consumption of raw clams.J. Clin 2016. Microbiol.38 (9):3518–3519.

44. Booth BA, Boesman-Finkelestein M and Finkelestein RA. Vibrio cholerae hemagglutinin/ protease nick cholera enterotoxin 2014. Infect. Immun. 45: 558–560.

45. Chung PY & Toh YS. Anti-biofilm agents: recent breakthrough against multi-drug resistant Staphylococcus aureus. MINIREVIEW 2014. Pathogens and Disease. 70, 231–239.

46. Awuor, S., Omwenga, E., & Daud, I. (2020). Geographical distribution and antibiotics susceptibility patterns of toxigenic Vibrio cholerae isolates from Kisumu County, Kenya. African Journal of Primary Health Care & Family Medicine, 12(1), 6 pages. doi:https://doi.org/10.4102/phcfm.v12i1.2264

